# Predicting cued and oddball visual search performance from neural representational similarity

**DOI:** 10.1101/2023.06.15.545065

**Authors:** Lu-Chun Yeh, Sushrut Thorat, Marius V. Peelen

**Author notes:** **Corresponding author:** Lu-Chun Yeh Mathematical Institute Justus Liebig University Gießen Arndtstraße 2 35392 Gießen, Germany. Conflict of interest statement. The authors declare no competing financial interests.

## Abstract

Capacity limitations in visual tasks can be observed when the number of task-related objects increases. An influential idea is that such capacity limitations are determined by competition at the neural level: two objects that are encoded by shared neural populations interfere more in behavior (e.g., visual search) than two objects encoded by separate neural populations. However, the neural representational similarity of objects varies across brain regions and across time, raising the question of where and when competition determines task performance. Furthermore, it is unclear whether the association between neural representational similarity and task performance is common or unique across tasks. Here, we used neural representational similarity derived from fMRI, MEG, and deep neural networks (DNN) to predict performance on two visual search tasks involving the same objects and requiring the same responses but differing in instructions: cued visual search and oddball visual search. Separate groups of human participants (both sexes) viewed the individual objects in neuroimaging experiments to establish the neural representational similarity between those objects. Results showed that performance on both search tasks could be predicted by neural representational similarity throughout the visual system (fMRI), from 80 msec after onset (MEG), and in all DNN layers. Stepwise regression analysis, however, revealed task-specific associations, with unique variability in oddball search performance predicted by early/posterior neural similarity, and unique variability in cued search task performance predicted by late/anterior neural similarity. These results reveal that capacity limitations in superficially similar visual search tasks may reflect competition at different stages of visual processing.

**Significance Statement:** Visual search for target objects is slowed down by the presence of distractors, but not all distractors are equally distracting – the more similar a distractor is to the target, the more it slows down search. Here, we used fMRI, MEG, and a deep neural network to reveal where, when, and how neural similarity between targets and distractors predicts visual search performance across two search tasks: oddball visual search (locating the different-looking object) and cued visual search (locating the cued object). Results also revealed brain regions, time points, and feature levels that predicted task-unique performance. These results provide a neural basis for similarity theories of visual search and show that this neural basis differs across visual search tasks.

## Introduction

Our daily-life environments contain multiple objects that are impossible to process all at once. According to the biased competition model of attention (Desimone & Duncan, 1995), this limitation is reflected in competition for neural representation in visual cortex, with attention being the mechanism that resolves this competition. A central tenet of the biased competition model is that competitive interactions are strongest between stimuli that activate neurons in nearby regions of visual cortex (Desimone, 1998). While most early studies focused on competition in visuotopically organized areas (e.g., Reynolds et al., 1999), more recent work has provided evidence that this principle extends to the representational space of objects in ventral temporal cortex (VTC; Reddy & Kanwisher, 2007; Kliger & Yovel, 2020; Bao & Tsao, 2020).

If the level of competition between objects is determined by their representational similarity in visual cortex, we would expect this to be reflected in behavior as well. Indeed, several studies have linked neural representational similarity in visual cortex to visual search efficiency (e.g., Sripati et al., 2010; Proklova et al., 2016), providing a link between neural similarity and stimulus similarity, as postulated in theories of visual search (Duncan & Humphreys, 1989). A more general link to behavior was demonstrated by a series of functional magnetic resonance imaging (fMRI) studies that related neural representational similarity to performance on multiple visual tasks (visual search, visual categorization, visual working memory, visual awareness; Cohen et al., 2014; 2015; 2017). In these studies, objects that evoked relatively similar fMRI activity patterns, when presented in isolation, were found to interfere more in these visual tasks. For example, participants were faster to search for a car among faces (relatively low neural similarity) than for a car among phones (relatively high neural similarity). For all tasks, this relationship was most strongly observed in ventral and lateral occipitotemporal cortex. Integrating the findings across studies, the authors concluded that the representational similarity of objects in visual cortex reflects “a stable architecture of object representation that is a primary bottleneck for many visual behaviors” (Cohen et al., 2017).

One interpretation of these findings is that different kinds of visual judgments involving the same set of objects would be limited by the objects’ representational similarity in the same parts of visual cortex. Alternatively, the relationship between neural representational similarity and visual task performance may differ across tasks. For example, tasks that rely more on low-level features could be constrained by the objects’ representational similarity in low-level visual cortex, whereas tasks relying on matching visual input to memory templates may be constrained by representational similarity in high-level visual cortex (Cohen et al., 2017). In the present study, we aimed to reveal the common and unique representational stages that constrain two superficially similar visual search tasks: cued visual search (Wolfe, 2020) and oddball visual search (Arun, 2012). As an example of these tasks, consider a fruit basket containing an apple and three oranges; on one occasion, we may search for the apple (*cued visual search*), while on another occasion, we may search for the different-looking object (*oddball visual search*). These tasks involve the same visual input and require the same response. However, previous research has shown that only performance on the cued visual search task is influenced by the categorical similarity of the objects (Belke et al., 2008, Moores et al., 2003; Telling et al., 2010; Yeh & Peelen, 2022). This raises the question of how performance on these tasks relates to the objects’ neural representational similarity in visual cortex, both spatially (fMRI) and temporally (magnetoencephalography; MEG).

Here, using the approach of Cohen et al. (2017), we related performance on cued and oddball visual search tasks to neural representational similarity using fMRI, MEG, and a deep neural network (DNN). This design allowed us to test where and when neural representational similarity predicts task performance that is common and/or unique to cued and oddball visual search, thereby informing theories of visual search.

## Materials and Methods

### Visual tasks

We examined competition at the behavioral level in three visual tasks involving the same object set (Figure 1A): cued visual search, oddball visual search, and a same/different task. Reaction time (RT) data were taken from previous studies, each involving a different group of participants (cued visual search task (N=16, 6 males and 10 females) and same/different task (N=24, 8 males and 16 females): Yeh & Peelen, 2022; oddball visual search task (N=18, 2 males and 16 females): Proklova et al., 2016). For all analyses, we took the reciprocal of RT (1/RT) as a measure of dissimilarity (i.e., faster responses indicate higher dissimilarity), following previous recommendations (Arun, 2012). The three tasks had different procedures but shared the same stimuli (Figure 1A). Eight animate and eight inanimate objects were used as stimuli, and each object had four exemplars (64 stimuli in total). All stimuli were grayscaled and were presented on a gray background.

**Figure 1.**
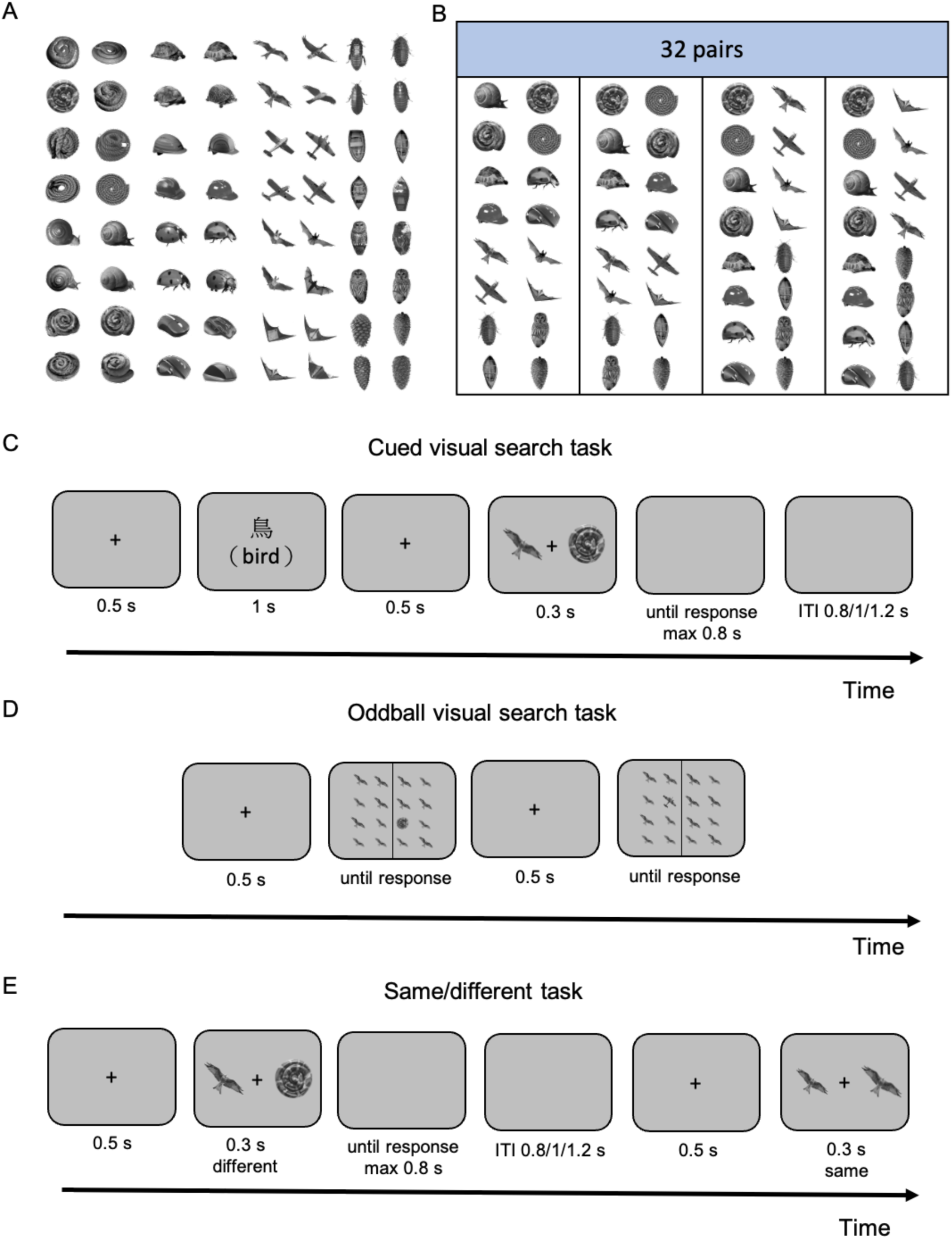
Stimuli and tasks. (A) All stimuli used in the experiments. (B) In the behavioral experiments, stimuli were paired based on categorical and perceptual similarity (Yeh & Peelen, 2022), for a total of 32 pairs. (C) Task procedure of the cued visual search task. Participants were asked to search for a predefined target (in Chinese, the cue in English was not shown in the real experiment) and press the button corresponding to the target location. (ITI: intertrial interval) (D) Task procedure of the oddball visual search task. Participants were asked to search for an odd stimulus and press the button corresponding to the target location. (E) Task procedure of the same/different task. Participants were asked to judge whether the two objects were the same or different.

In the cued visual search task (Figure 1C), participants were instructed to search for a predefined target (e.g., searching for a bird), and indicate the target location (left or right side of the search array). The search array contained a target and a distractor, and each object was set as the target with equal frequency. In the oddball visual search task (Figure 1D), participants indicated the location of the oddball object. In each oddball search display, 16 objects were presented in a 4-by-4 grid, consisting of one oddball target and 15 identical distractors (at slightly different sizes). Finally, in the same/different task (Figure 1E), participants indicated whether the two objects were the same or different. The task consisted of two conditions, same-object and different-object conditions. In the same-object condition, the display consisted of two identical objects. The size of the objects was varied, with one object being 90%, 100%, or 110% as large as the other, but this was irrelevant for the task. In the different-object condition, the display consisted of two different objects at their original size. For more details, see the methods of Experiment 1 and 2 in Yeh & Peelen (2022) and Experiment 1 in Proklova et al. (2016).

We added the same/different task based on previous research showing that variations in RT across object pairs in the same/different task are highly correlated with variations in RT in the oddball task (Jacob & Arun, 2020). We replicated this finding in our data. Specifically, we computed the correlation between performance on the oddball and same/different tasks for corresponding object pairs using a split-half correlation analysis. We first randomly assigned participants to one of two groups and then computed the correlation between the object pair means of the two datasets both within and between tasks. We repeated this procedure 100 times and computed the average of the outputs. We found that performance on the oddball visual search task was closely related to performance on the same/different task, with the between-task correlation (r = 0.85) being close to the within-task correlations (same/different: r = 0.90; oddball: r = 0.97). This confirms that the same/different task and the oddball visual search task provided similar measures of pairwise task performance. To examine whether same/different task performance was more similar to oddball than cued search task performance, we correlated individual same/different data with the group-averaged oddball and cued search task data. Then, we used the Fisher z-transformed correlation coefficient to run the paired t-test (two-tailed). The result showed that the correlation between the same/different and oddball task data (𝑍_*r*_ = 0.65) was significantly higher than the correlation between the same/different and cued task data (𝑍_*r*_ = 0.57) (*p* =.002). We also ran the reverse analyses, correlating individual oddball/cued task data with the averaged same/different task data and then ran an independent t-test (two-tailed). The result again showed that the correlation between the same/different and oddball task data (𝑍_*r*_ = 1.07) was significantly higher than the correlation between the same/different and cued task data (𝑍_*r*_ = 0.81) (*p* <.001).

### Neural measurements and representational dissimilarity

Neural responses evoked by each individual object were recorded with fMRI and MEG. These data were taken from previous studies (fMRI (N=18, 7 males and 11 females): Proklova et al., 2016; MEG (N=29, 16 males and 13 females): Proklova et al., 2019). During those recordings, only one object was presented at a time and no search task was performed.

The representational dissimilarity of the fMRI data was computed using the approach of Proklova et al. (2016), restricted to the 32 pairs used in the behavioral experiments (Figure 1B). Briefly, following Proklova et al. (2016), we computed the correlation between the beta values of the objects in each pair across the voxels of spherical searchlight spheres (100 voxels each). Then, the representational similarity values were transformed into dissimilarity values by subtracting the correlation values from 1.

The representational dissimilarity of the MEG data was computed using pairwise decoding analyses for each time point (10 msec) for each participant, following Proklova et al. (2019). Neural dissimilarity is reflected by the decoding accuracy: the higher the decoding accuracy, the greater the neural dissimilarity. We used the 32 pairs of neural dissimilarity results of magnetometers (Proklova et al. (2019) reported similar results for magnetometers and gradiometers). We combined data across the two experiments reported in Proklova et al. (2019).

Finally, we established the representational dissimilarity of the object set in AlexNet (Krizhevsky et al., 2017), a feedforward DNN for object classification. We first obtained activations elicited by the stimuli (individual objects) at each layer of AlexNet. For each relevant pair of stimuli, at each layer, the similarity between the activation patterns was computed using correlation. Then, the representational similarity values were transformed into dissimilarity values by subtracting the correlation values from 1. The schematic of the analysis approach is illustrated in Figure 2.

**Figure 2.**
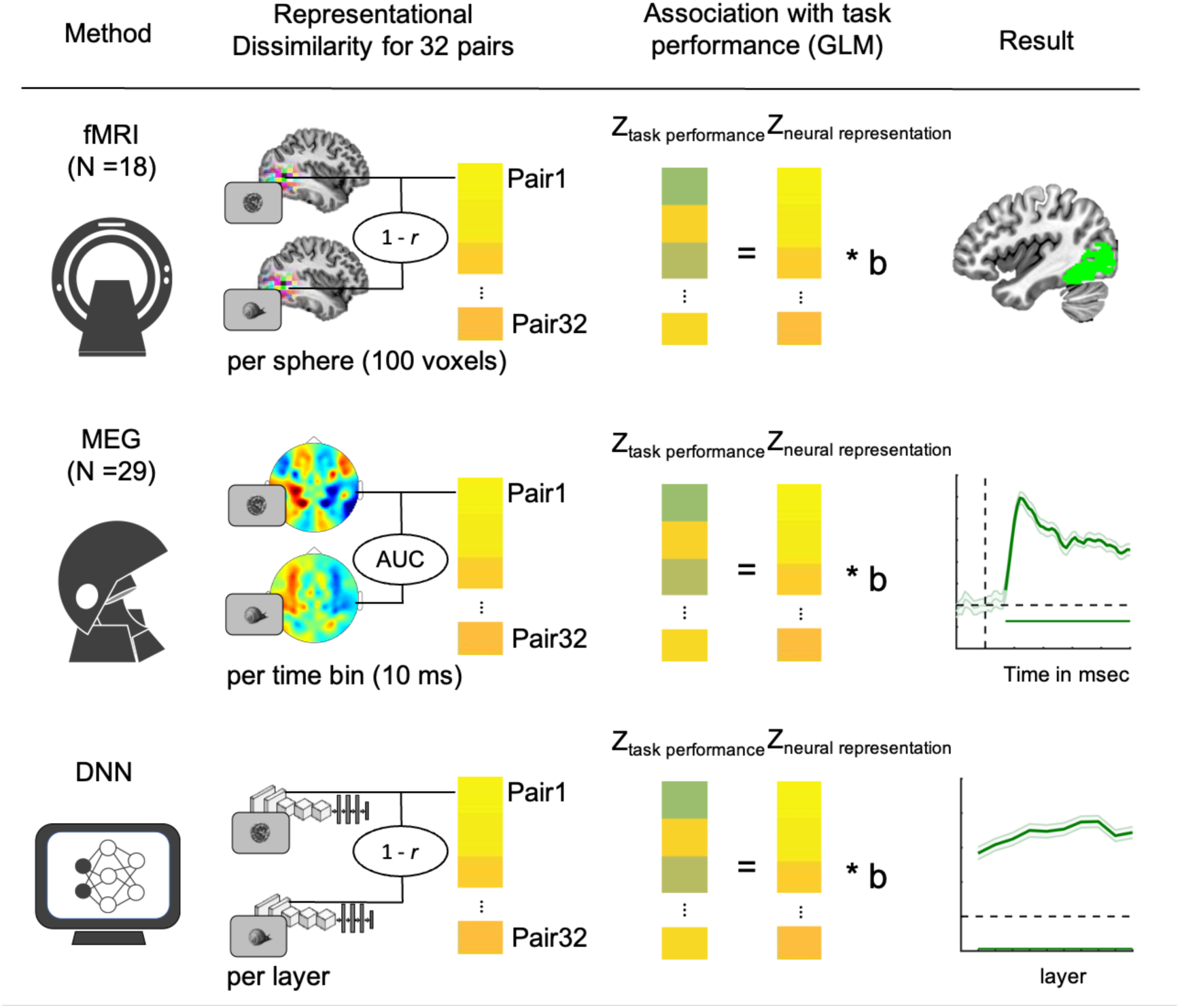
Schematic of analysis approach. Neural activity to individual objects was measured with fMRI, MEG, and DNN. These activity measures were used to compute the neural representational dissimilarity for 32 pairs of stimuli that were presented in the visual tasks. Finally, neural representational dissimilarity was used to predict task performance using regression analysis. This analysis can show where, when, and in which layer neural representational dissimilarity predicts task performance.

### Relating neural representational dissimilarity to task performance

We used linear regression analysis to relate pairwise neural representational dissimilarity to pairwise task performance. We used the z-transformed neural dissimilarity to predict the z-transformed task performance (1/RT). For fMRI and MEG analyses, we used the dissimilarity values of 32 pairs from each participant to predict the group-average task performance in random-effects statistical analyses across the participants in the neuroimaging experiments. This way, the statistical analyses tested for consistency across the participants of the fMRI and MEG studies, with the averaged behavioral data as to-be-predicted values. Please note that this approach is more conservative than the reverse approach of averaging the neural data and testing across individual behavioral participant data, considering that the neural data are likely to be much more variable than the behavioral data. Indeed, the split-half within-task correlations for the behavioral data were very high (same/different: r = 0.90; oddball: r = 0.97; cued: r =0.94). Analyses that used the reverse approach - averaging the neuroimaging data and testing across the behavioral participants - gave qualitatively similar and statistically more significant results than those reported here, confirming that the behavioral data were more consistent across participants than the neuroimaging data. Here, we report the results across the participants of the neuroimaging experiments to address our research question concerning the neural correlates of two types of visual search behavior. Note, however, that this approach was not possible for the DNN analysis, for which there were no participants, such that for that analysis we used the dissimilarity values obtained from AlexNet to predict task performance for each participant in the behavioral tasks (again using random-effects analyses). This resulted in beta maps (fMRI), beta timelines (MEG), and betas for each layer (DNN), reflecting the associations between neural representational dissimilarity and performance on each of the three tasks (Figure 2).

Furthermore, to explore unique associations between representational dissimilarity and task performance, we ran two-step regression analyses to (1) compare the cued visual search task with the oddball visual search task and (2) compare the cued visual search task with the same/different task. To calculate betas reflecting task-unique associations, we first ran a linear regression analysis using one task performance (e.g., cued visual search) to predict the other task performance (e.g., oddball visual search). Then, we ran another linear regression analysis using the *z*-transformed neural dissimilarity values to predict the *z*-transformed *residual* of the first regression analysis. Overall, we obtained four beta maps (fMRI), beta timelines (MEG), and betas for each layer (DNN) for three tasks. Two are for the cued visual search task when removing the effect of the oddball visual search task and the same/different task, one is for the oddball visual search task when removing the effect of the cued visual search task, and one is for the same/different task when removing the effect of the cued visual search task.

### Statistical tests

To test the significance of the beta values, we used one-sample t-tests against zero and controlled the type I error for multiple comparisons. For the fMRI analyses, we used a voxel-level threshold of *p* < .001 (uncorrected) and a cluster-level threshold of *p* < .05 (FWE corrected) to control type I error for the whole brain. Clusters were only reported when the cluster size was larger than 50 voxels. For fMRI MNI y-coordinate, MEG, and DNN analyses, the statistical significance of correlations was tested using cluster-based nonparametric permutation one-sample *t*-tests against zero to determine the window of significant correlation, controlling for multiple comparisons (Maris & Oostenveld, 2007). The permutation test was with null distribution created from 1000 Monte Carlo random partitions using a cluster correction for type I error (p <.05, one-tailed).

## Results

### fMRI: Spatial associations between neural representational dissimilarity and task performance

To test for brain regions in which neural representational dissimilarity predicted task performance, we performed single regression analyses using a whole-brain searchlight approach (see Materials and Methods). For each sphere, neural representational dissimilarity (based on responses evoked by objects presented in isolation) was used to predict pairwise task performance (Figure 2), separately for the three tasks. Results showed that performance on all three tasks was predicted by representational dissimilarity in both low- and high-level visual cortex (Figure 3A; Table 1), in line with previous reports (Proklova et al., 2016; Cohen et al., 2017). This was confirmed by an analysis that averaged beta values across x- and z-coordinates, with significantly positive beta values in both posterior and anterior visual cortex regions (Figure 4, solid lines). These results thus reveal considerable overlap in the brain regions that predicted performance on each task.

**Figure 3.**
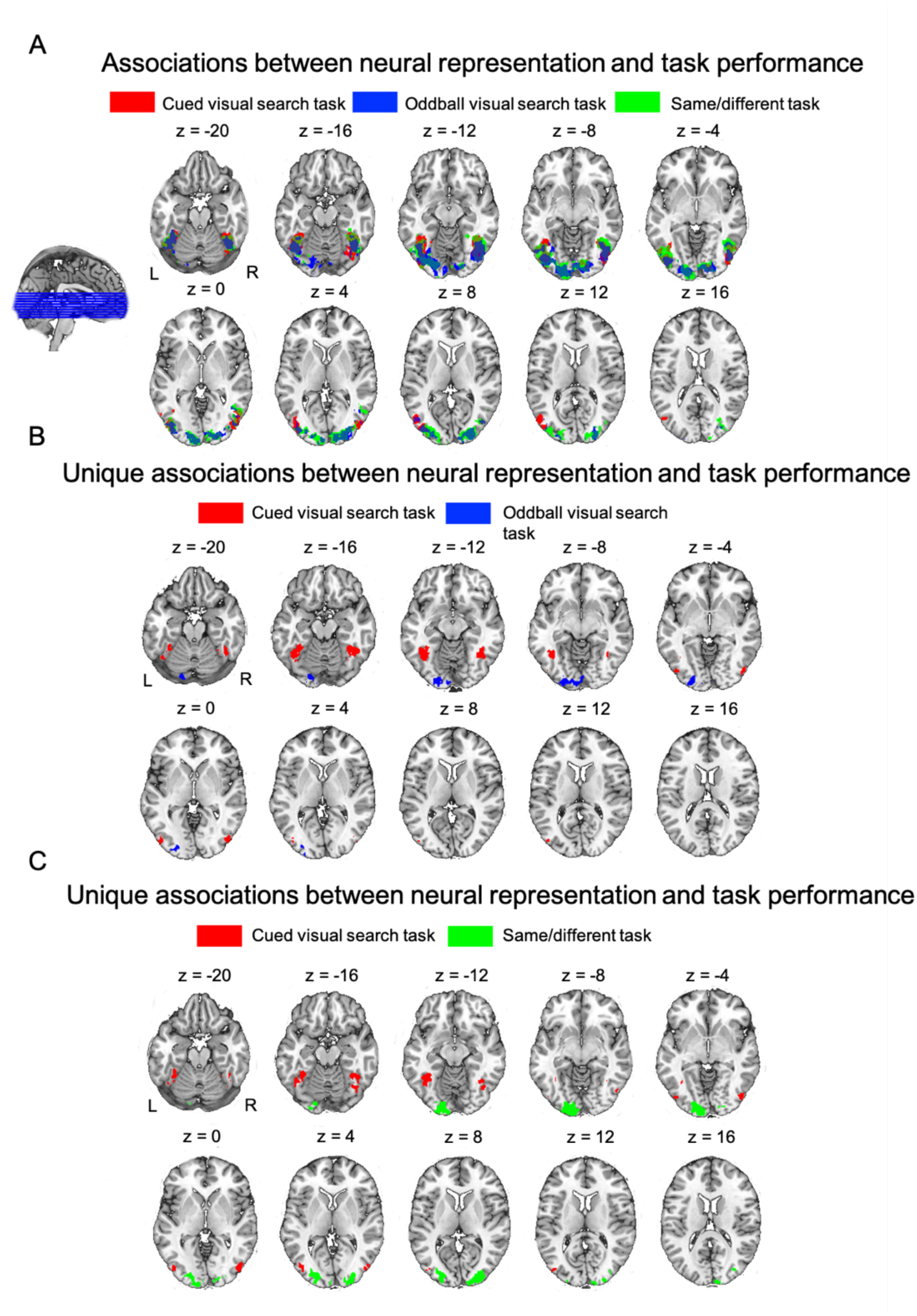
Whole-brain fMRI searchlight maps relating neural representational dissimilarity to task performance. A) Clusters in which neural representational dissimilarity predicted task performance, separately for each task. B) Clusters in which neural representational dissimilarity predicted performance unique to the cued visual search task (red) and the oddball visual search task (blue). C) Clusters in which neural representational dissimilarity predicted performance unique to the cued visual search task (red) and the same/different task (green).

**Figure 4.**
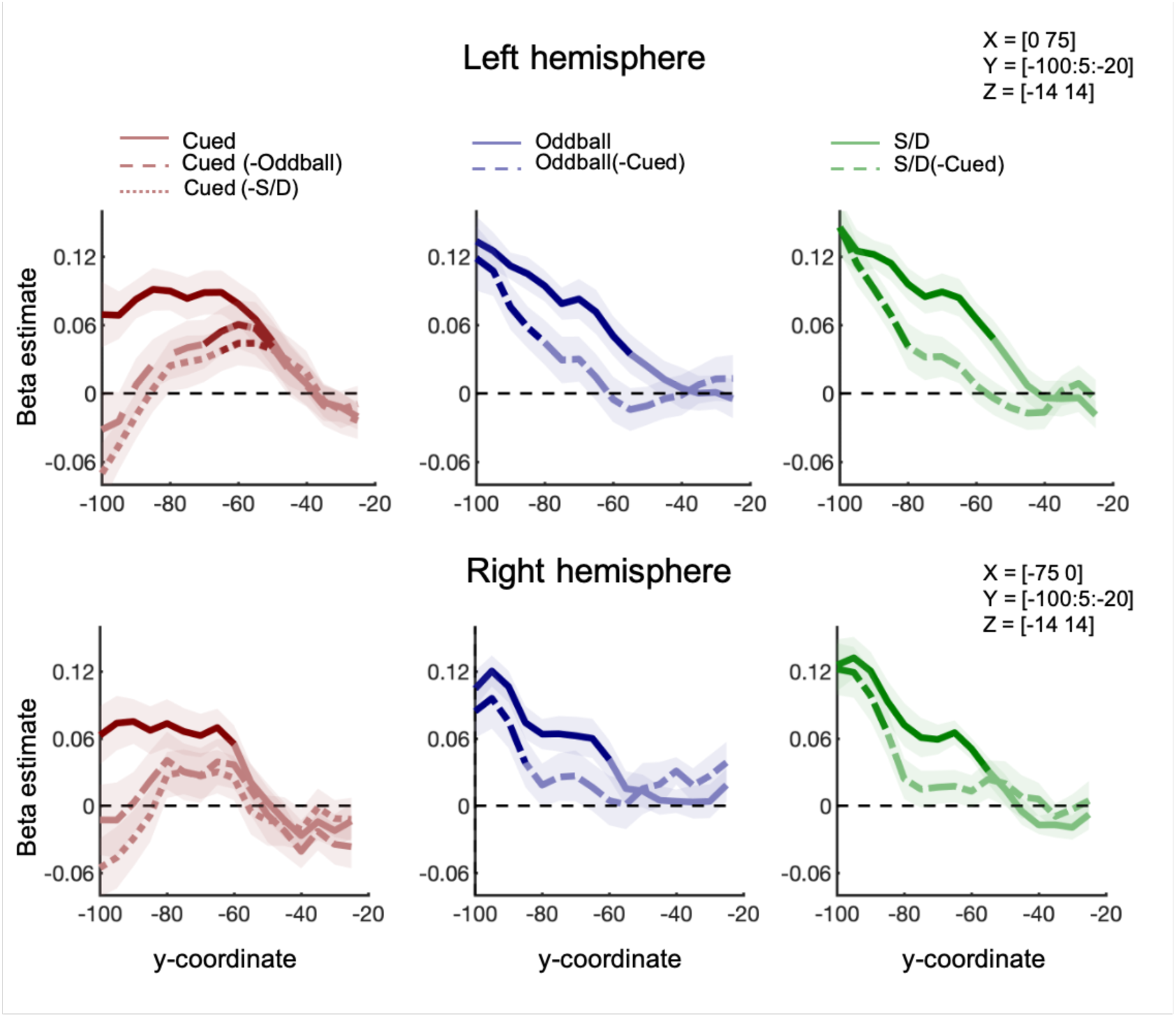
fMRI results across the visual hierarchy. Graphs show the average beta estimate along the y-coordinate axis for left (top panel) and right (bottom panel) hemispheres. The means were computed for every 5 mm in the MNI y-coordinate (averaging across the x and z coordinates). Solid lines show results from analyses relating neural representational dissimilarity to performance on each task separately. Dashed lines show results from analyses relating neural representational dissimilarity to performance on one task, after regressing out performance on another task (stated after the “-” symbol; e.g., “Cued (-Oddball)” refers to cued visual task performance that excluded the covariance with oddball task performance). Significant clusters (cluster-based permutation tests, *p* < .05, one-tailed) are highlighted in bold.

**Table 1.**
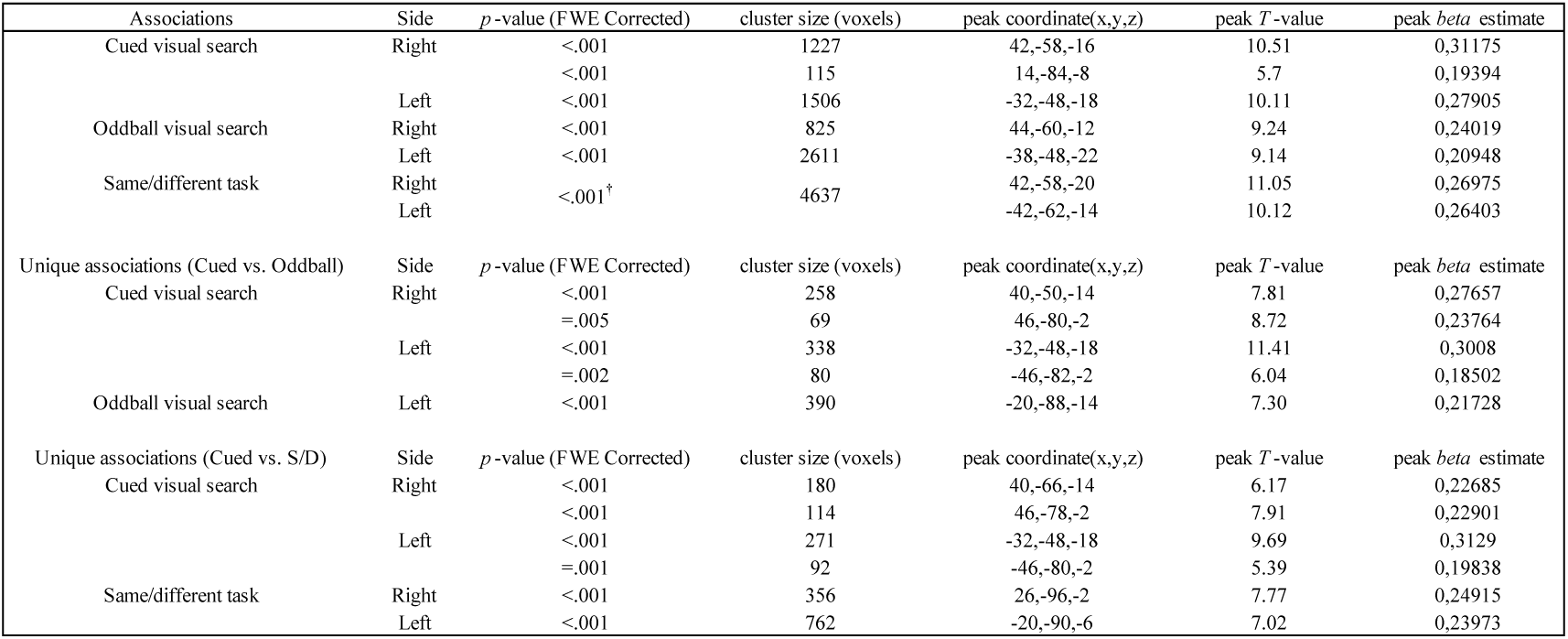
All significant clusters in the whole-brain fMRI analyses. Peak coordinates are in MNI space. ^†^The significant cluster for the same/different task includes both right and left hemispheres.

Next, we tested for task-specific associations between neural representational dissimilarity and task performance by first regressing out performance on the oddball visual search task from performance on the cued visual search task (and vice versa). The same analysis was repeated for the same/different task and the cued visual search task. Results revealed several brain regions in which neural representational dissimilarity predicted variability in task performance that was unique to one of the tasks (Figure 3B; Table 1). Specifically, unique variability in cued visual search performance was predicted by relatively anterior regions in visual cortex, overlapping the fusiform gyrus, while unique variability in oddball visual search performance was predicted by more posterior regions. This result was replicated when including the same/different task instead of the oddball task (Figure 3C; Table 1), showing that these unique associations did not simply reflect differences in the displays of cued and oddball visual search tasks (Figure 1). The posterior-anterior difference was also clearly observed in the y-coordinate analysis (Figure 4), with the unique prediction of the cued visual search task strongest at y=-60 and the unique prediction of the oddball visual search task strongest at the most posterior parts of visual cortex (y=-100). These results thus reveal that performance on two highly similar visual search tasks are uniquely predicted by neural representational dissimilarity in different brain regions.

### MEG: Temporal associations between neural representational dissimilarity and task performance

Recent MEG studies have shown that patterns of activity across MEG sensors can be used to decode object properties in a time-sensitive manner (Grootswagers et al., 2017; Robinson et al., 2023), revealing the representational dynamics of visual processing (Carlson et al., 2013; Cichy et al., 2014). Using this approach, we tested *when* neural representational dissimilarity predicts visual search performance. The regression analysis linking neural representational dissimilarity to task performance was otherwise identical to that performed on the fMRI data (Figure 2). Results showed that performance on all three tasks was predicted by neural representational dissimilarity from 80 msec after stimulus onset (cluster-based *p* <.001; Figure 5). The analyses testing for task-specific associations revealed unique associations between neural representational dissimilarity and task performance of the oddball visual search task (cluster-based *p* <.001) and the same/different task (cluster-based *p* <.001) starting from around 80 ms after stimulus onset (Figure 5). Unlike the fMRI results, no unique association was found between neural representational dissimilarity and cued visual search task performance. These results provide a clear temporal association between neural representational dissimilarity and task performance, and show that the unique association between neural representational dissimilarity and cued visual search task performance that we observed in fMRI was not captured by MEG activity patterns.

**Figure 5.**
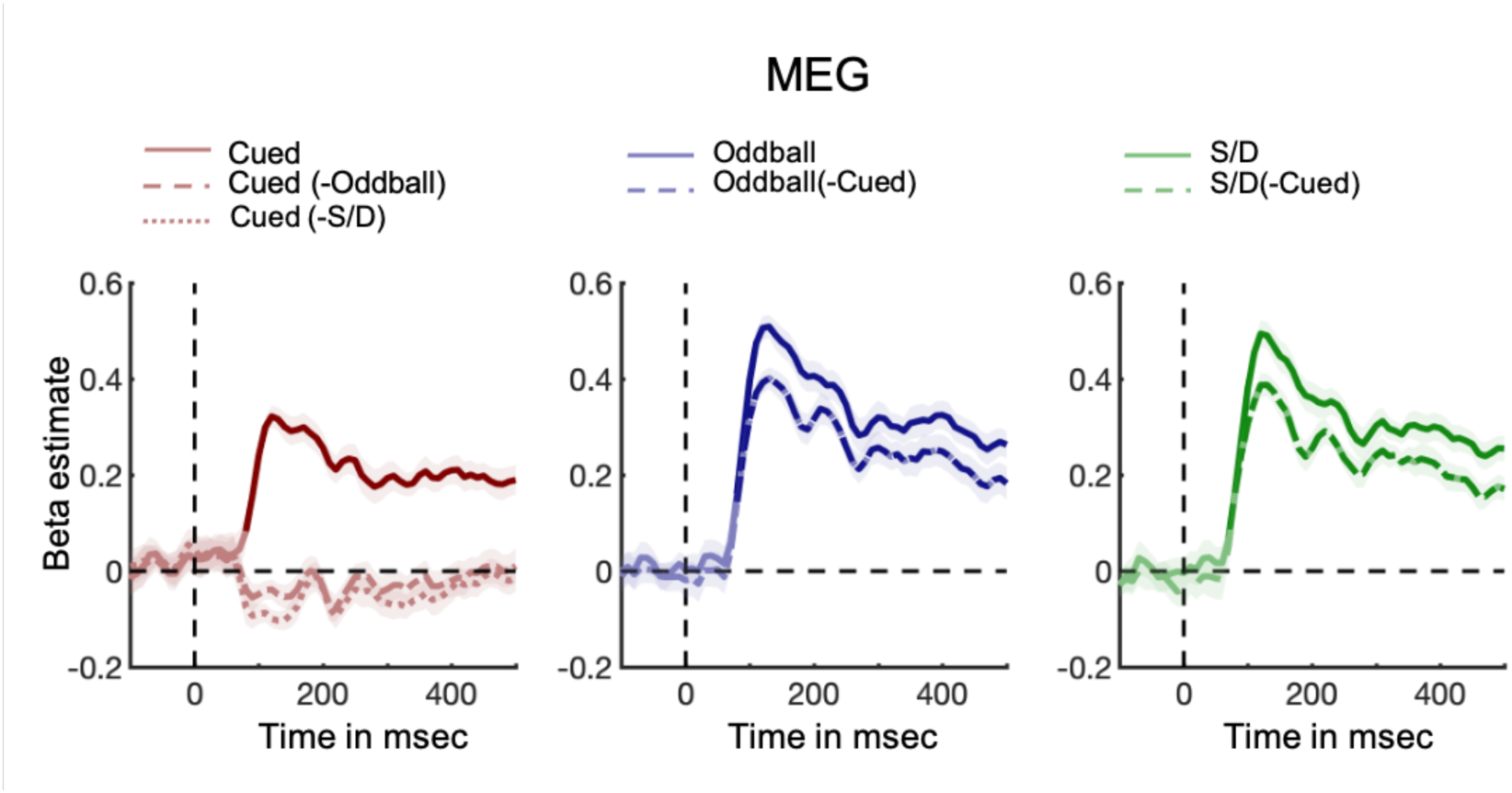
MEG results showing temporal associations between neural representational dissimilarity and task performance. MEG results of the regression analyses using neural representational dissimilarity to predict cued visual search task performance (left panel), oddball visual search performance (middle panel), and same/different task performance (right panel). Solid lines show results from analyses relating neural representational dissimilarity to performance on each task separately. Dashed lines show results from analyses relating neural representational dissimilarity to performance on one task, after regressing out performance on another task (stated after the “-” symbol; e.g., “Cued (-Oddball)” refers to cued visual task performance that excluded the covariance with oddball task performance). Significant clusters (cluster-based permutation tests, *p* < .05, one-tailed) are highlighted in bold.

### DNN: Layer-wise associations between representational dissimilarity and task performance

Finally, we used a feedforward DNN model (AlexNet) to explore which layers of the hierarchical network predicted task performance. We followed the regression approach we used in the fMRI/MEG analysis but now using the representational dissimilarity computed from the patterns of activation in DNN layers (Figure 2). Results showed that performance on all three tasks was predicted by the representational dissimilarity of all ten selected layers (cluster-based *p* <.001; Figure 6): the Input layer, the five convolutional layers (Conv1, Conv2, Conv3, Conv4, and Conv5), the three fully-connected layers (FC6, FC7, FC8), and the softmax layer on the readout (SM). Here, we also found task-specific associations. The cued visual search task performance was predicted by representational dissimilarity in the deeper layers, including FC6, FC7, FC8, and SM (cluster-based *p* =.005) after regressing out oddball visual search performance, and was predicted by representational dissimilarity in FC7, FC8, and SM (cluster-based *p* =.0036) after regressing out same/different task performance (Figure 6). By contrast, oddball visual search and same/different task performance were predicted by all layers after regressing out cued visual search performance, and contrary to the cued visual search, most strongly with the earlier layers (cluster-based *p* <.001).

**Figure 6.**
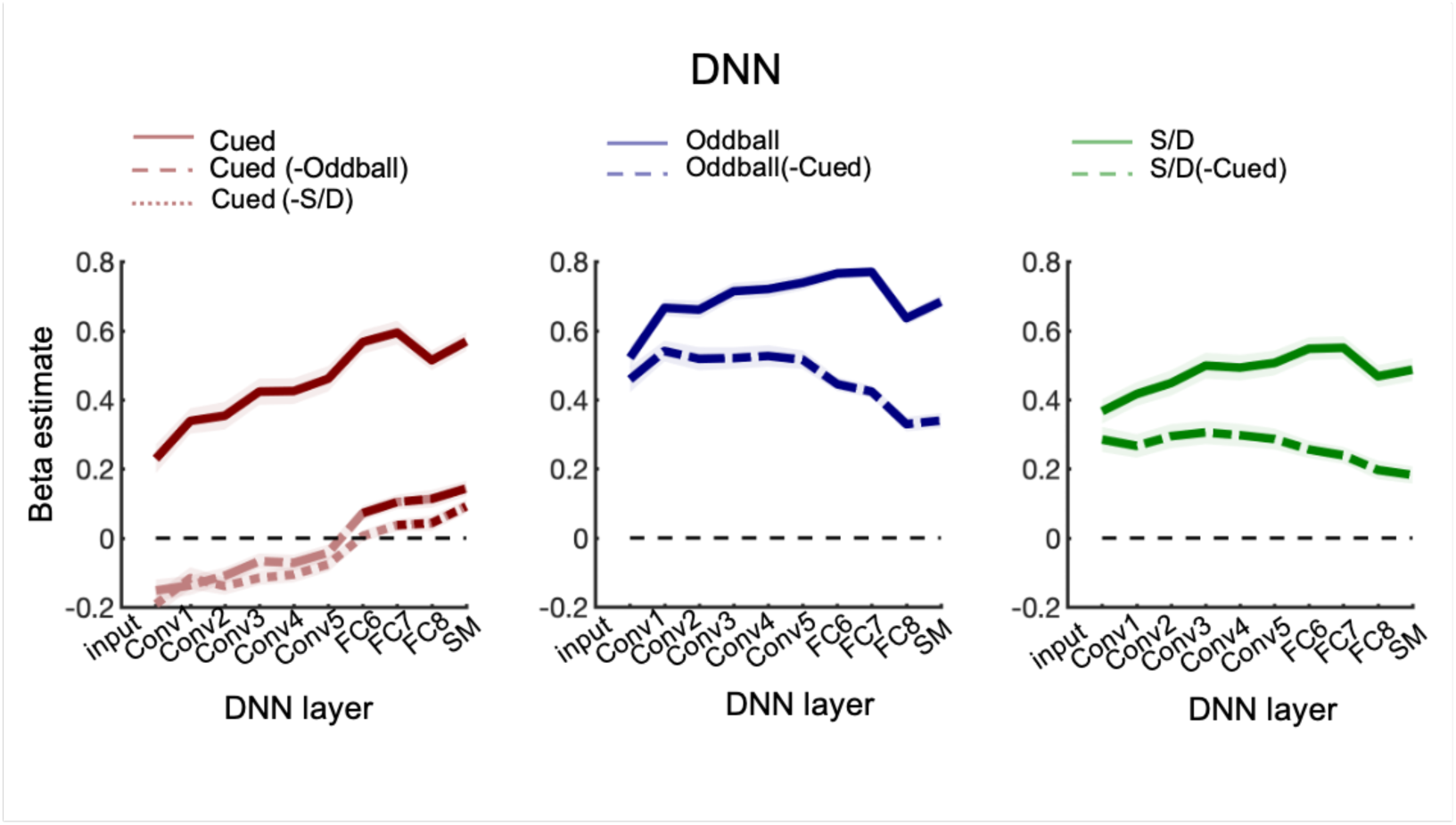
DNN results showing associations between representational dissimilarity and task performance. DNN results of the regression analyses using representational dissimilarity to predict the cued visual search performance (left panel), oddball visual search performance (middle panel), and same/different task performance (right panel). Solid lines show results from analyses relating representational dissimilarity to performance on each task separately. Dashed lines show results from analyses relating neural representational dissimilarity to performance on one task, after regressing out performance on another task (stated after the “-” symbol; e.g., “Cued (-Oddball)” refers to cued visual task performance that excluded the covariance with oddball task performance). Significant clusters (cluster-based permutation tests, *p* < .05, one-tailed) are highlighted in bold.

## Discussion

In this study, we tested whether, where, and when the pairwise neural representational similarity of objects predicts visual search performance (RT) involving these object pairs. By using fMRI, MEG, and a DNN we established a detailed spatiotemporal association between neural representational similarity and visual search performance. Results showed that visual search performance was predicted by neural representational similarity in both high- and low-level regions of the visual cortex (fMRI), from 80 msec after onset (MEG), and in all DNN layers. When analyzed separately, similar brain-behavior associations were observed for a cued visual search task and an oddball visual search task. This shared component is in line with previous results showing that performance on different tasks can be predicted by the same stable neural similarity structure (Cohen et al., 2014; 2015; 2017).

Importantly, however, using stepwise regression analysis, we could relate these task-unique components of performance to neural representational similarity as well. Results showed that neural representational similarity in regions of high-level visual cortex (fMRI) and the late layers of a DNN predicted the unique component of the cued visual search task. By contrast, neural representational similarity in regions of low-level visual cortex (fMRI) and the early layers of a DNN predicted the unique component of the oddball visual search task. These results provide evidence that performance on superficially similar visual search tasks, involving the same stimuli and response, is nonetheless constrained by neural similarity at different stages of visual processing.

The finding that search performance is predicted by neural similarity in visual cortex is in line with theories that explain visual search efficiency in terms of stimulus similarity (Duncan & Humphreys, 1989). Specifically, search performance has been proposed to be determined by a combination of target-distractor and distractor-distractor similarity (the pairwise approach in the current study only considers target-distractor similarity). The similarity theory of visual search used the term similarity to refer to similarity along multiple dimensions, with the relevant dimension depending on the experimental stimuli (e.g., color, shape). For example, in an experiment where color was the only manipulated dimension, color similarity would determine search efficiency (Duncan & Humphreys, 1989). In the current experiment, we investigated search for familiar real-world objects that differed along multiple feature dimensions (e.g., texture, shape). Interestingly, the association between neural similarity and visual search was observed throughout the visual cortex (fMRI), across a long time window (MEG), and at different hierarchical levels of visual feature representation (DNN). These findings therefore suggest that the similarity that predicts visual search performance for familiar objects is multidimensional, likely consisting of a combination of low- and high-level features.

At the same time, our results show that the similarity dimension that determines search performance depends on the type of search task, even for identical stimuli. Specifically, similarity at higher levels of visual cortex and in later layers of a neural network predicted the unique component of cued visual search. These unique associations may reflect features that are diagnostic of the category (animate/inanimate) of the objects, in line with previous behavioral and EEG studies that found that cued visual search is influenced by categorical similarity (Belke et al., 2008, Moores et al., 2003; Telling et al., 2010; Yeh & Peelen, 2022). Furthermore, in a previous fMRI study (Proklova et al., 2016), we showed that the categorical similarity of the object set was represented in high-level visual cortex regions that overlap with the regions that predicted the unique cued visual search performance in the current study. Finally, the representational similarity in later layers of AlexNet have been shown to follow a categorical organization (Khaligh-Razavi & Kriegeskorte, 2014).

One possibility is that the high-level component observed for the cued visual search task reflects an additional template-matching stage, comparing visual input to a target template held in working memory (Cohen et al., 2017; Wolfe, 2021; Eimer, 2014). In this scenario, the cued visual search task could be equivalent to the oddball visual search task plus an additional template-matching stage. Interestingly, however, our results also revealed unique brain-behavior associations for the oddball task, particularly in earlier stages of visual processing. These results suggest that different stimulus dimensions can be flexibly emphasized to support different visual search tasks on the same stimulus set. This is in line with a growing literature demonstrating the flexible adjustment of top-down attentional templates, for example based on expected distractor characteristics (Lee & Geng, 2020; Lerebourg et al., 2023, Boettcher et al., 2020).

The current MEG results showed that representational similarity from 80 ms after stimulus onset reliably predicted visual search performance. By contrast, however, the unique component of cued visual search performance was not reflected in the representational structure of MEG at any time point, even though unique associations were observed in comparable fMRI and DNN analyses. These results indicate that while MEG sensor patterns are highly sensitive to the low-/mid-level object features that predict visual search performance in general, they are not sensitive to the high-level (e.g., categorical) features that predict the unique aspect of cued visual search performance. This finding is in line with a previous study in which MEG patterns did not discriminate between perceptually matched object categories, while fMRI patterns did (Proklova et al., 2019). The absence of categorical information in MEG patterns may appear in contrast to studies that have successfully decoded object category using MEG (e.g., animacy; Carlson et al., 2013; Cichy et al., 2014). However, these studies did not control for perceptual similarity, such that decoding could reflect mid-level feature differences between categories (e.g., curvature). Proklova et al. (2019) proposed several explanations for the discrepancy between MEG and fMRI decoding of object category, including differences in spatial resolution and temporal averaging. Another possible reason for the absence of categorical information in MEG patterns is that MEG is not sensitive to deep sources. However, we similarly observed no significant association between representational similarity and the unique aspect of cued visual search performance in an EEG dataset (data not shown), with EEG being more sensitive to deep sources than MEG. The lack of sensitivity of M/EEG to high-level object features means that it remains an open question when during visual processing neural representational similarity predicts the unique aspect of cued visual search.

Our approach has several limitations. First, our analyses were done across different groups of participants, focusing on commonalities across groups. This approach prevents analyses that link representational structure to behavioral performance at the individual-subject level (Charest et al., 2014). Second, we used a stimulus set in which half the object pairs were matched for outline shape and that included a clear categorical distinction. Higher-level dimensions may become particularly relevant in visual search for objects that are perceptually similar (Yeh & Peelen, 2022). Therefore, future studies would need to replicate our results with a larger and more diverse stimulus set. Third, our search tasks were highly simplified, involving only one distractor object, fixed presentation locations, and no scene context. By contrast, real-world search typically includes many distractors, variable locations, and meaningful context. Finally, the approach of associating stable neural architecture to visual search performance is, by definition, indirect, such that a positive association between neural similarity and task performance in a particular brain region (or time point) does not prove that that brain region causally determines task performance.

## Conclusion

In summary, our results reveal a strong association between neural representational similarity and visual search performance: when the neural representation of a target object was relatively dissimilar to the neural representation of a distractor object, visual search was relatively fast. The finding that unique performance on two superficially similar visual search tasks was predicted by neural similarity in different visual cortex regions and in different DNN layers shows that neural constraints on visual task performance are, to some extent, task specific. Capacity limitations in superficially similar visual search tasks may thus reflect competition at different stages of visual processing.

## Acknowledgements

This project has received funding from the European Research Council (ERC) under the European Union’s Horizon 2020 research and innovation programme (grant agreement No. 725970).

## Notes

### Competing Interest Statement

The authors have declared no competing interest.

### Summary of Updates

Methods and Discussion updated.

## References

Arun, S. P. (2012). Turning visual search time on its head. Vision Research, 74(1), 86–92.

Bao, P., & Tsao, D. Y. (2018). Representation of multiple objects in macaque category-selective areas. Nature communications, 9(1), 1–16.

Belke, E., Humphreys, G. W., Watson, D. G., Meyer, A. S., & Telling, A. L. (2008). Top-down effects of semantic knowledge in visual search are modulated by cognitive but not perceptual load. Perception & Psychophysics, 70(8), 1444–1458.

Boettcher, S. E., van Ede, F., & Nobre, A. C. (2020). Functional biases in attentional templates from associative memory. Journal of Vision, 20(13), 7.1–10.

Carlson, T., Tovar, D. A., Alink, A., & Kriegeskorte, N. (2013). Representational dynamics of object vision: the first 1000 ms. Journal of vision, 13(10), 1.1–19.

Charest, I., Kievit, R. A., Schmitz, T. W., Deca, D., & Kriegeskorte, N. (2014). Unique semantic space in the brain of each beholder predicts perceived similarity. Proceedings of the National Academy of Sciences, 111(40), 14565–14570.

Cichy, R. M., Pantazis, D., & Oliva, A. (2014). Resolving human object recognition in space and time. Nature neuroscience, 17(3), 455–462.

Cohen, M. A., Alvarez, G. A., Nakayama, K., & Konkle, T. (2017). Visual search for object categories is predicted by the representational architecture of high-level visual cortex. Journal of Neurophysiology, 117(1), 388–402.

Cohen, M. A., Konkle, T., Rhee, J. Y., Nakayama, K., & Alvarez, G. A. (2014). Processing multiple visual objects is limited by overlap in neural channels. Proceedings of the National Academy of Sciences of the United States of America, 111(24), 8955–8960.

Cohen, M. A., Nakayama, K., Konkle, T., Stantić, M., & Alvarez, G. A. (2015). Visual awareness is limited by the representational architecture of the visual system. Journal of Cognitive Neuroscience, 27(11), 2240–2252.

Desimone, R., & Duncan, J. (1995). Neural mechanisms of selective visual attention. Annual Review of Neuroscience, 33(18), 193–222.

Desimone, R. (1998). Visual attention mediated by biased competition in extrastriate visual cortex. Philosophical Transactions of the Royal Society of London. Series B: Biological Sciences, 353(1373), 1245–1255.

Duncan, J., & Humphreys, G. W. (1989). Visual search and stimulus similarity. Psychological review, 96(3), 433–458.

Duncan, J., Humphreys, G., & Ward, R. (1997). Competitive brain activity in visual attention. Current opinion in neurobiology, 7(2), 255–261.

Eimer, M. (2014). The neural basis of attentional control in visual search. Trends in cognitive sciences, 18(10), 526–535.

Grootswagers, T., Wardle, S. G., & Carlson, T. A. (2017). Decoding dynamic brain patterns from evoked responses: A tutorial on multivariate pattern analysis applied to time series neuroimaging data. Journal of cognitive neuroscience, 29(4), 677–697.

Jacob, G., & Arun, S. P. (2020). How the forest interacts with the trees: Multiscale shape integration explains global and local processing. Journal of Vision, 20(10), 20.1-21.

Kliger, L., & Yovel, G. (2020). The functional organization of high-level visual cortex determines the representation of complex visual stimuli. Journal of Neuroscience, 40(39), 7545–7558.

Khaligh-Razavi, S. M., & Kriegeskorte, N. (2014). Deep supervised, but not unsupervised, models may explain IT cortical representation. PLoS computational biology, 10(11), e1003915.

Krizhevsky, A., Sutskever, I., & Hinton, G. E. (2017). ImageNet Classification with Deep Convolutional Neural Networks. Communications of the ACM, 60(6), 84–90.

Lee, J., & Geng, J. J. (2020). Flexible weighting of target features based on distractor context. *Attention, Perception*, & Psychophysics, 82, 739–751.

Lerebourg, M., de Lange, F. P., & Peelen, M. V. (2023). Expected Distractor Context biases the Attentional Template for Target Shapes. Journal of Experimental Psychology: Human Perception and Performance, 49(9), 1236–1255.

Maris, E., & Oostenveld, R. (2007). Nonparametric statistical testing of EEG-and MEG-data. Journal of neuroscience methods, 164(1), 177–190.

Moores, E., Laiti, L., & Chelazzi, L. (2003). Associative knowledge controls deployment of visual selective attention. Nature neuroscience, 6(2), 182–189.

Proklova, D., Kaiser, D., & Peelen, M. V. (2016). Disentangling representations of object shape and object category in human visual cortex: The animate–inanimate distinction. Journal of cognitive neuroscience, 28(5), 680–692.

Proklova, D., Kaiser, D., & Peelen, M. V. (2019). MEG sensor patterns reflect perceptual but not categorical similarity of animate and inanimate objects. Neuroimage, 193, 167–177.

Robinson, A. K., Quek, G. L., & Carlson, T. A. (2023). Visual Representations: Insights from Neural Decoding. Annual Review of Vision Science, 9,2.1–2.23.

Reddy, L., & Kanwisher, N. (2007). Category selectivity in the ventral visual pathway confers robustness to clutter and diverted attention. Current Biology, 17(23), 2067–2072.

Reynolds, J. H., Chelazzi, L., & Desimone, R. (1999). Competitive mechanisms subserve attention in macaque areas V2 and V4. Journal of Neuroscience, 19(5), 1736–1753.

Sripati, A. P., & Olson, C. R. (2010). Global image dissimilarity in macaque inferotemporal cortex predicts human visual search efficiency. Journal of Neuroscience, 30(4), 1258–1269.

Telling, A. L., Kumar, S., Meyer, A. S., & Humphreys, G. W. (2010). Electrophysiological Evidence of Semantic Interference in Visual Search. Journal of Cognitive Neuroscience, 22(10), 2212–2225.

Wolfe, J. M. (2020). Visual search: How do we find what we are looking for? Annual review of vision science, 6, 539–562.

Wolfe, J. M. (2021). Guided Search 6.0: An updated model of visual search. Psychonomic Bulletin & Review, 28(4), 1060–1092.

Yeh, L.-C., & Peelen, M. V. (2022). The time course of categorical and perceptual similarity effects in visual search. Journal of Experimental Psychology: Human Perception and Performance, 48(10), 1069–1082.

